# Orientation and color tuning of the human visual gamma rhythm

**DOI:** 10.1101/2021.04.23.441193

**Authors:** Ye Li, William Bosking, Michael S. Beauchamp, Sameer A. Sheth, Daniel Yoshor, Eleonora Bartoli, Brett L. Foster

**Author notes:** <u>Corresponding author:</u> Brett L. Foster, Eleonora Bartoli.

## Abstract

Narrowband gamma oscillations (NBG: ∼20-60Hz) in visual cortex reflect rhythmic fluctuations in population activity generated by underlying circuits tuned for stimulus location, orientation, and color. Consequently, the amplitude and frequency of induced NBG activity is highly sensitive to these stimulus features. For example, in the non-human primate, NBG displays biases in orientation and color tuning at the population level. Such biases may relate to recent reports describing the large-scale organization of single-cell orientation and color tuning in visual cortex, thus providing a potential bridge between measurements made at different scales. Similar biases in NBG population tuning have been predicted to exist in the human visual cortex, but this has yet to be fully examined. Using intracranial recordings from human visual cortex, we investigated the tuning of NBG to orientation and color, both independently and in conjunction. NBG was shown to display a cardinal orientation bias (horizontal) and also an end- and mid-spectral color bias (red/blue and green). When jointly probed, the cardinal bias for orientation was attenuated and an end-spectral preference for red and blue predominated. These data both elaborate on the close, yet complex, link between the population dynamics driving NBG oscillations and known feature selectivity biases in visual cortex, adding to a growing set of stimulus dependencies associated with the genesis of NBG. Together, these two factors may provide a fruitful testing ground for examining multi-scale models of brain activity, and impose new constraints on the functional significance of the visual gamma rhythm.

**Significance Statement:** Oscillations in electrophysiological activity occur in visual cortex in response to stimuli that strongly drive the orientation or color selectivity of visual neurons. The significance of this induced ‘gamma rhythm’ to brain function remains unclear. Answering this question requires understanding how and why some stimuli can reliably generate gamma activity while others do not. We examined how different orientations and colors independently and jointly modulate gamma oscillations in the human brain. Our data show gamma oscillations are greatest for certain orientations and colors that reflect known biases in visual cortex. Such findings complicate the functional significance of gamma activity, but open new avenues for linking circuits to population dynamics in visual cortex.

**Classification:** Neuroscience

## Introduction

Within visual cortex, high frequency fluctuations in electrophysiological activity can be induced by visual stimulation. This induced ‘gamma’ range (∼20-200 Hz) activity is composed of at least two distinct spectral responses: asynchronous broadband- or ‘high-’ gamma (BBG; ∼70-200 Hz) and oscillatory narrow-band gamma (NBG; ∼20-60 Hz) (1, 2). NBG has been the subject of detailed experimental (3, 4) and computational (5) study, leading to a proposed role in synchronizing functional brain dynamics important for perception and cognition (6). While these efforts have shown that early reports of visual NBG activity can be replicated across species (7), a continued debate still surrounds its functional significance (8). In recent years, this debate has focused on the stimulus dependence of visually induced NBG (9, 10). Despite long standing evidence (11, 12), it is now clear that standard grating or Gabor patch stimuli used in early studies of visual cortex were particularly suited at inducing NBG. In contrast, more complex stimuli (e.g. natural images) often fail to evoke robust NBG responses (2, 12, 13). While grating-like stimuli are particularly effective at driving NBG, the magnitude and peak frequency of this response is sensitive to grating attributes such as size (14, 15), contrast (2, 16) and spatial frequency (14, 17), adding an additional layer of stimulus dependence. Most recently, it has also been shown that visual NBG can be induced by uniform color stimuli, with a predominate preference for long wavelength hues (i.e. red/orange) (2, 18). Interestingly, gamma responses induced by grating or uniform color stimuli can be contextually modulated by discontinuities between the receptive field and surround (19, 20). Together, this dependence on structural and chromatic stimulus features can account for much of the difficulty that has been observed in predicting which complex, and in particular natural, stimuli will evoke NBG oscillations (2, 9, 10, 13, 21, 22). It is therefore critical to piece together the diverse evidence of stimulus-dependencies to better determine the correspondence between visual features, cortical tuning preferences and the genesis of visual gamma oscillations. Through linking gamma rhythms to underlying visual area circuit properties, we can advance our understanding of their functional significance in vision.

What factors can account for both the structural and chromatic dependence of visual gamma? While orientation and color are both processed in early visual cortex, they have historically been viewed as having modest overlap, suggesting different populations drive these gamma effects. However, it has long been known that this is a simplified dichotomy, as neurons in visual cortex show complex tuning properties to orientation and color stimuli (23, 24). Building on these observations, recent imaging work in the non-human primate has provided clear evidence for joint orientation and color tuning by neurons in early visual cortex (25). This joint tuning suggests activation of overlapping circuits may account for the structural and chromatic tuning of visual gamma (21). To date, no prior studies have examined the effect of combined color and orientation information on the emergence of gamma oscillations in visual cortex. However, recent work has shown visual gamma is sensitive to incongruous center-surround attributes for both color or orientation stimuli (19, 26, 27). Therefore, visual gamma rhythms can be induced and enhanced by specific stimulus configurations uniformly driving features strongly represented in the locally recorded population – such that NBG provides a partial readout of underlying large-scale circuit properties, like population tuning bias for some stimulus features (21). If correct, the visual gamma rhythm should display a dependence and tuning profile similar to prior observations of such population tuning activity in early visual cortex. Indeed, evidence from the non-human primate suggests that visual gamma displays putative cardinal orientation tuning (17) – a bias in population tuning also observed in human visual cortex through neuroimaging (28, 29). As noted above, recent work has also shown chromatic tuning of visual gamma, which shows some correspondence to previously reported end-spectral biases (i.e. stronger responses to red and blue) in early visual cortex (30). Crucially, orientation tuning of gamma has yet to be systematically examined with direct recordings in the human brain, and no previous study has addressed how this tuning relates to color nor the joint tuning of orientation and color. For example, does the human visual gamma rhythm also display population biases in orientation and color, and do these properties interact when both features are visually driven?

To address these questions, we performed direct intracranial recordings from early visual cortex in the human brain, in order to quantify orientation, color and joint orientation-color tuning of the human visual gamma rhythm. Specifically, we focused on changes in NBG activity, indicative of rhythmic activity, in contrast to the commonly observed broadband gamma (BBG) or ‘high-gamma’ activity (2). Across four experiments, we observed NBG to reveal cardinal orientation bias, as well as both end- and mid-spectral color biases. Interestingly, when jointly driving orientation and color together, cardinal orientation and mid-spectral color tuning were attenuated while end-spectral bias was maintained. These findings suggest a striking relationship between visual gamma tuning and recent observations about the distribution of orientation and color tuning within visual cortex (25). This tight link between functional circuit properties and visual gamma tuning provides a fruitful avenue for empirically and computationally bridging physical scales of organization in the brain, while providing important constraints and new directions for the functional significance of gamma oscillations in vision and cognition more broadly.

## Results

### Visual Electrode Identification

We employed electrocorticography (ECoG) and stereo-electroencephalography (sEEG) recordings from human visual cortex in 7 subjects being invasively monitored for the potential surgical treatment of refractory epilepsy (Table 1). Across subjects, we obtained 121 electrodes within anatomically defined occipital cortex (Figure 1A, see details in Table S1). To exclude electrodes lacking robust visual responses from our analysis, we further applied a functional criterion to define visual responsiveness based on the presence of a visual evoked potential (VEP) in response to a grating stimulus. As shown in Figure 1B, clear voltage deflections are observed after stimulus onset in VEP electrodes. In addition, we assigned each electrode to putative functional subdivisions of visual cortex using a probabilistic atlas (V1-4, see Methods). Of all the electrodes within visual cortex, ∼76% displayed a VEP (92/121). Of all VEP electrodes, 50 were within V1, 18 were within V2/V3/V4, and 24 were outside of V1-4 (see Table S1).

**Table 1.**
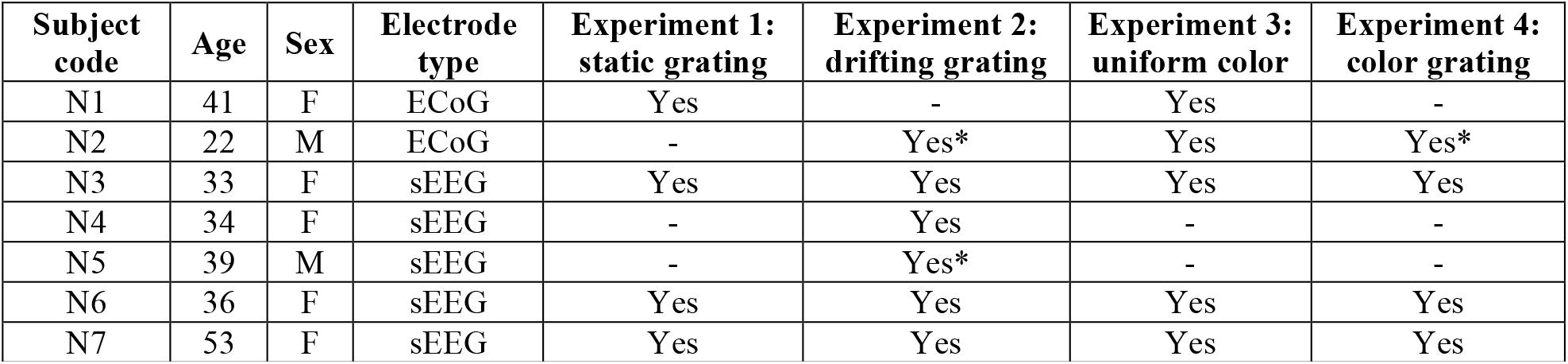
Subject information. For experiments 2 and 4, the asterisks (*) denotes that a subset of the orientations were recorded (4 in place of 8 orientations).

**Figure 1.**
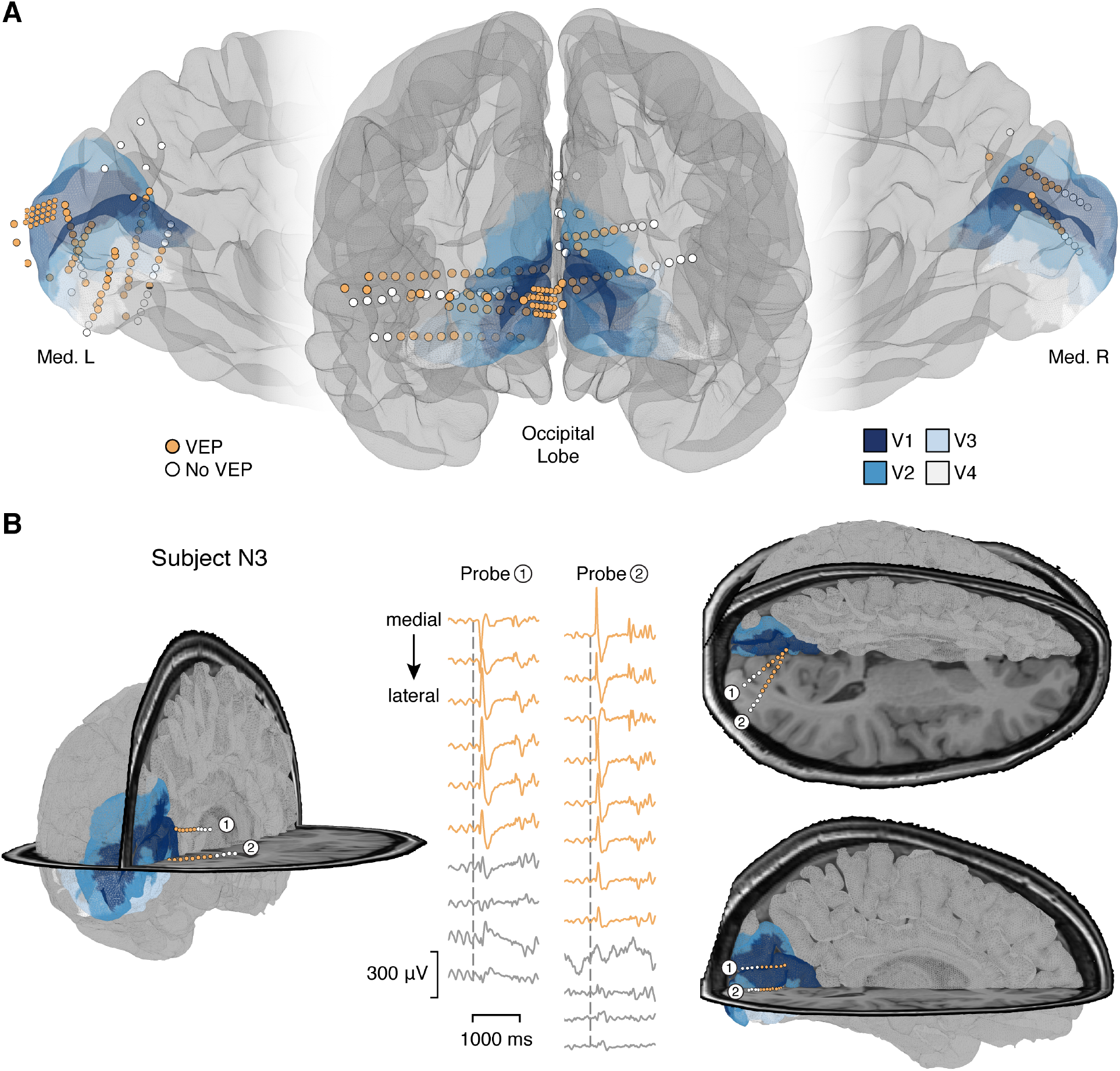
Recording Sites within Visual Cortex. **(A)** Anatomical location of electrodes from all subjects participating in the current study on a standard cortical surface (see Methods). VEP electrodes are shown in orange, non-VEP electrodes are shown in white. The V1-4 areas from vcAtlas are mapped on the standard cortical surface (see method). **(B)** Anatomical location of electrodes from an sEEG case (subject N3). Inset shows the mean voltage traces for each electrode of two probes shown in the figure. Orange indicates sites and traces of VEP identified electrodes.

### Peak Frequency of Induced NBG is increased for Drifting vs. Static Grating Contrast

A large and growing literature shows visual grating stimuli to be particularly effective at inducing NBG activity (1, 2, 13). Consistent with this literature, we recently reported a strong positive correlation between grating contrast level and induced NBG peak frequency in human visual cortex (2). In the current study, we sought to employ drifting grating stimuli, which are particularly effective at driving robust responses in early visual cortex.

To provide a bridge with our prior observations, we first replicated the grating contrast dependence of NBG responses for static stimuli. In addition, as we previously utilized invasively implanted surface ECoG electrodes to measure visual NBG responses, the present replication served to ensure similar effects were captured with sEEG depth electrodes. We therefore examined the amplitude and peak frequency of induced NBG as a function of contrast for both static and drifting grating stimuli. In Experiment 1 (static grating task), four subjects (N1, N3, N6-7) were presented with full-screen static grayscale gratings at three contrast levels (20%, 50%, and 100%; spatial frequency 1 cycle/degree), for a 500 ms duration. In Experiment 2 (drifting grating task), six subjects (N2-7) were presented with full-screen drifting grayscale gratings at three contrast levels (20%, 50%, and 100%; spatial frequency 1 cycle/degree; velocity 2 degree/sec), for a 500 ms duration.

Consistent with prior studies, we observed clear induced oscillatory responses in the raw voltage during both static and drifting grating presentation (Figure 2A). As mentioned above, NBG amplitude and peak frequency are sensitive to the contrast level of static gratings (2, 16, 31). To replicate this finding in the current study, we computed the normalized amplitude spectra of the 250-500 ms post-stimulus time window for the static grating task. As shown for a representative subject in Figure 2B, the peak frequency of NBG monotonically increases for higher grating contrast levels (see Figure S1 for the group averaged spectra). Similar results were found in the drifting grating task when calculating the normalized amplitude spectra for each contrast condition by averaging across other conditions (i.e., orientations and motion directions). Group averaged spectrograms for both tasks show a similarly sustained NBG response during stimulus presentation (Figure 2C), in addition to large transient BBG responses at stimulus onset, closely matching prior studies (2, 16). Indeed, both tasks showed qualitatively similar increases in NBG amplitude across contrast levels (Figure 2D). However, as shown in Figure 2E, drifting gratings induced a systematically higher NBG peak frequency compared to static gratings, while still maintaining a monotonic increase with higher contrast levels.

**Figure 2.**
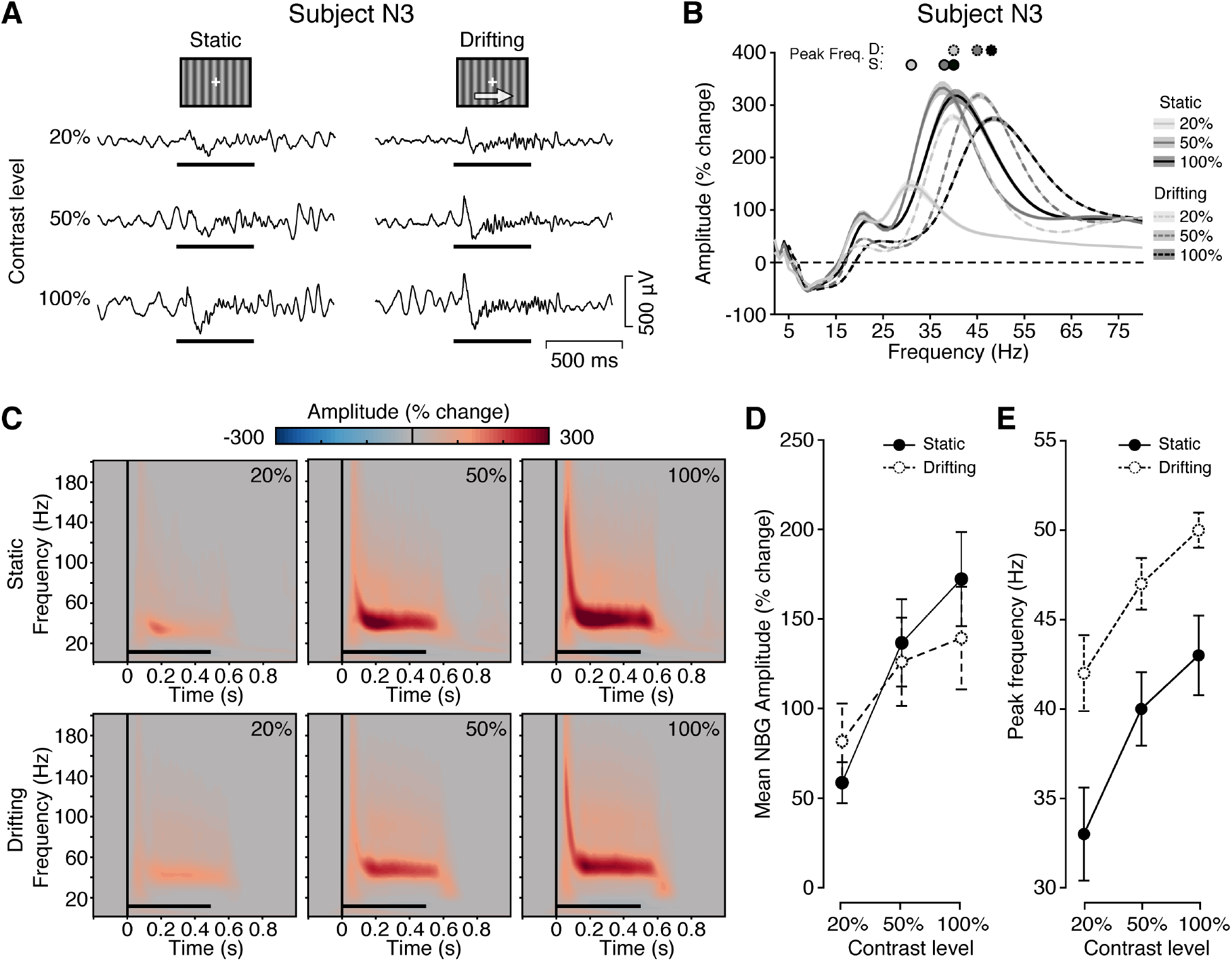
Experiment 1 and 2: Spectral Response to Static and Drifting Grating Stimuli. **(A)** Example single trial voltage responses (from subject N3) for each contrast level in Experiment 1 and 2. Grating stimuli were presented for 500 ms (black horizontal line), with a random inter-stimulus interval (ISI) of 1.5-2.0 s. Drifting grating motion was in one of two orthogonal directions for each trial (directions balanced across trials). **(B)** Normalized amplitude spectra (from all VEP electrodes of subject N3) for each contrast level in Experiment 1 and 2 averaged from the 250-500 ms post-stimulus time window. Filled circles indicate peak amplitude frequency (D: drifting grating; S: static grating). Shading reflects standard error. The peak frequency of NBG monotonically increases for higher grating contrast levels in both tasks. **(C)** Group average spectrograms for each grating contrast level in Experiment 1 (from subject N1, N3, N6, and N7) and 2 (from subject N2-7). Color maps indicate percentage change in amplitude relative to the pre-stimulus period (−500 to 0 ms). Black horizontal line indicates stimulus presentation. **(D)** Group average mean NBG amplitude in Experiment 1 and 2 (average of subjects N3, N6, and N7, who completed both experiments). Error bar indicates +/- 1 standard error. **(E)** Group average NBG peak frequency in Experiment 1 and 2 (average of subjects N3, N6, and N7, who completed both experiments). Error bar indicates +/- 1 standard error. While the monotonic increase with higher contrast maintained, drifting gratings enhanced the peak frequency of NBG for three contrast levels.

Quantitatively, NBG amplitude increased on average ∼57.5 % for each contrast level increment in the static grating task and ∼29.4 % in the drifting grating task, as shown in Table 2. The peak frequency of induced NBG increased on average ∼3.7 Hz for each contrast level increment in the both the static and the drifting grating tasks (Table 2). However, as noted above, the drifting grating task produced a NBG peak frequency that was on average 7.7 Hz higher than the static task (averaged across subjects who completed both Experiment 1 and 2: N3, N6-7). We employed mixed effect modeling to evaluate the statistical significance of the effect of contrast (see Methods and summary in Table 3). Grating stimuli contrast levels significantly modulated NBG amplitude across both experiments (static grating: *p* < 0.0001; drifting grating: *p* < 0.0001). Similar results were found when modeling the peak frequency of NBG (static grating: *p* < 0.0001; drifting grating: *p* < 0.0001). Together these data show parametric differences in NBG induced by static vs. drifting gratings, which are equally well captured by ECoG and sEEG recordings.

**Table 2.**
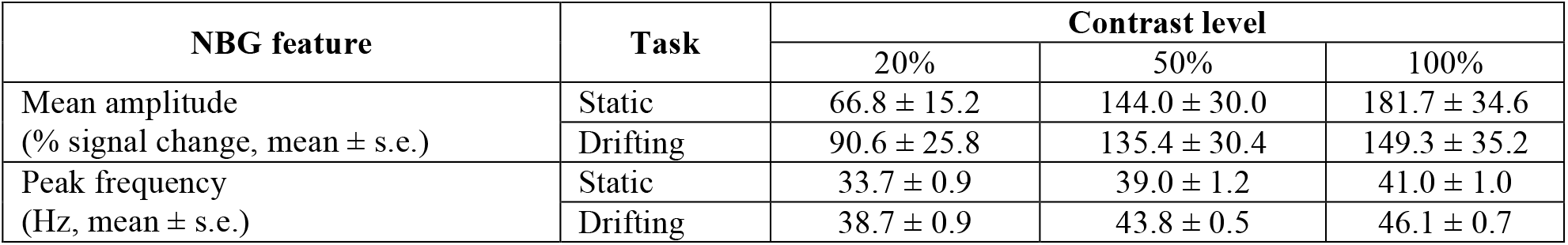
Descriptive statistics of mean amplitude and peak frequency of induced NBG in Experiment 1 and 2. Group averaged NBG amplitudes and peak frequencies for achromatic grating tasks (static/drifting) across contrast levels, using all subjects that participated in each task.

| NBG feature | Task | Contrast level |  |  |
| --- | --- | --- | --- | --- |
|  |  | 20% | 50% | 100% |
| Mean amplitude<br>(% signal change, mean ± s.e.) | Static | 66.8 ± 15.2 | 144.0 ± 30.0 | 181.7 ± 34.6 |
| | Drifting | 90.6 ± 25.8 | 135.4 ± 30.4 | 149.3 ± 35.2 |
| Peak frequency<br>(Hz, mean ± s.e.) | Static | 33.7 ± 0.9 | 39.0 ± 1.2 | 41.0 ± 1.0 |
| | Drifting | 38.7 ± 0.9 | 43.8 ± 0.5 | 46.1 ± 0.7 |

**Table 3.** Coefficients of fixed parameters in full models and complex models. *β*_*intercept*_ captures the 20% contrast level mean in each model. *β*_50%_ captures the mean difference between 50% and 20% contrast. *β*_100%_ captures the mean difference between 100% and 20% contrast. *β*_*motion*_ captures the mean difference between the drifting and static grating tasks in the complex models.

| Model components |  | Dataset | Coefficients (mean + s.e.) |  |  |  |
| --- | --- | --- | --- | --- | --- | --- |
| Dependent variable | Fixed effects | | $\beta_{intercept}$ | $\beta_{50\%}$ | $\beta_{100\%}$ | ($\beta_{motion}$) |
| Mean log amplitude<br>(% signal change) | Contrast | Static | 4.3 ± 0.1 | 0.6 ± 0.02 | 0.8 ± 0.02 | |
| | | Drifting | 4.6 ± 0.2 | 0.3 ± 0.01 | 0.4 ± 0.01 | |
| | Contrast + motion | Two tasks | 4.5 ± 0.2 | 0.3 ± 0.01 | 0.4 ± 0.01 | 0.05 ± 0.01 |
| Peak frequency (Hz) | Contrast | Static | 33.8 ± 1.1 | 5.2 ± 0.4 | 7.2 ± 0.4 | |
| | | Drifting | 38.5 ± 0.4 | 5.3 ± 0.2 | 7.8 ± 0.2 | |
| | Contrast + motion | Two tasks | 33.2 ± 0.3 | 5.3 ± 0.2 | 7.7 ± 0.2 | 5.4 ± 0.2 |

While NBG responses are sensitive to the contrast level in both experiments, the peak frequency is further increased by the drifting grating stimuli (i.e., grating motion). To quantify the grating motion effect on NBG responses, we combined data from both experiments and assessed the significance of grating motion effect (drifting vs. static). Both the mean log amplitude and peak frequency of induced NBG were significantly higher in the drifting grating experiment (see Table 3, coefficients *β*_*motion*_; NBG amplitude: *p* < 0.005; NBG peak frequency: *p* < 0.0001). Considering the mean log amplitude, the marginal increase in amplitude for drifting gratings occurred for the lowest contrast condition (20%), whereas at higher contrast levels the effect was reversed, with a tendency toward higher amplitude in response to static gratings. This was captured by a significant interaction between grating motion and contrast level (*p* < 0.0001). Considering peak frequency, post-hoc comparisons (see Methods) confirmed that the NBG peak frequencies were significantly higher in the drifting grating task at each contrast level (all *p* < 0.05 Bonferroni corrected, *p*_*corr*_). Indeed, the average NBG peak frequency recorded in response to static gratings at 100% contrast was indistinguishable from that of drifting gratings at 20% contrast (*p* = 0.49). The above analysis of the drifting grating data averaged responses across trials with different grating orientation and motion directions. Similar results for contrast were found when we considering each motion direction and cardinal orientation separately (see control analysis in Method and Table S2).

### NBG Displays Cardinal Orientation Selectivity

In quantifying the impact of grating contrast and motion, we collapsed data across different grating orientations. However, intracortical studies in the non-human primate have reported orientation tuning of NBG (14, 32). Interestingly, unlike the broad range of orientation preferences observed for spiking and multi-unit activity, NBG recorded simultaneously via the local field potential (LFP) shows a spatially consistent orientation preference (14, 17, 32, 33). Whereby, NBG is turned predominantly to cardinal orientations, in particularly 90° (17, 33). While such a ‘cardinal bias’ has been reported in human visual cortex with neuroimaging (28), the specific orientation tuning properties of NBG have been under examined in the human brain (34). Therefore, the invasive recordings of the current study provide a unique opportunity to directly test orientation tuning of NBG in the human brain. We next examined NBG responses across orientations (from 0° to 157.5°, with a 22.5° interval) from the drifting grating task (Experiment 2).

Examination of raw voltage data suggested subtle but consistent modulation of induced oscillations across orientations. As shown in Figure 3A, for an example subject (N7) and electrode, raw voltage responses showed induced oscillations that were more pronounced for the 90° orientation (horizontal). Averaged spectrograms of the same electrode revealed induced oscillations reflected sustained NBG responses, which were relatively higher in amplitude at 90° (Figure 3B). Examination of the normalized amplitude spectra for each orientation (percentage change of the 250-500 ms post-stimulus time window, Figure 3C) confirmed NBG amplitude was largest for 90° orientation (polar plot insert shows the NBG preference around 90°, with a circular mean of 80.6°). As shown in Figure 3D, mean normalized NBG amplitude across electrodes showed a tuning preference for 90°, consistently across grating contrast levels.

**Figure 3.**
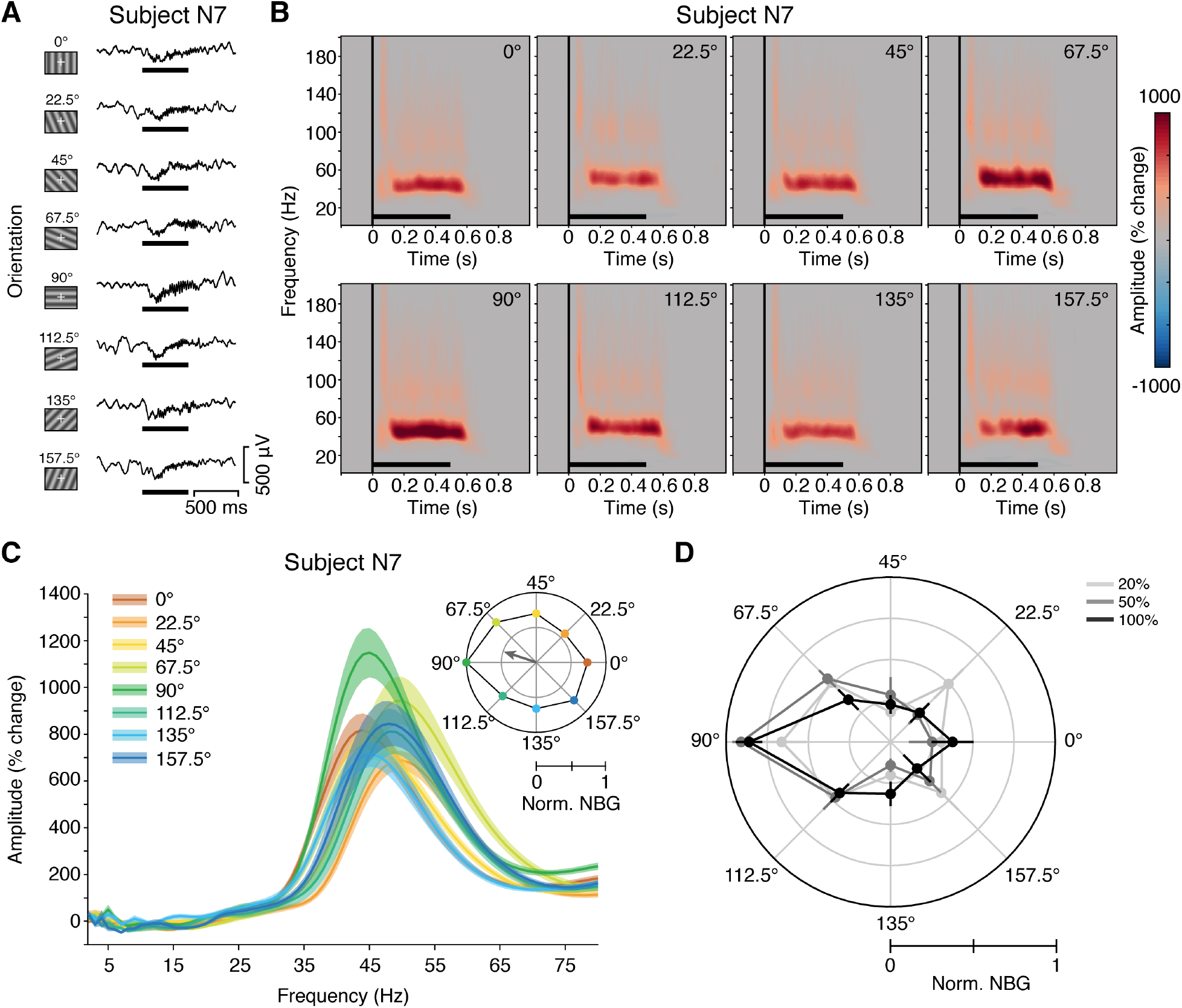
Experiment 2: Spectral Response to Drifting Grating Orientation. **(A)** Example single trial voltage responses (from subject N7) for each orientation at 100% contrast level in Experiment 2. Grating stimuli were presented for 500 ms (black horizontal line), with a random inter-stimulus interval (ISI) of 1.5-2.0 s. **(B)** Spectrograms for each orientation from all 100% contrast trials from the same electrode in (A). Color maps indicate percentage change in amplitude relative to the pre-stimulus period (−500 to 0 ms). Black horizontal line indicates stimulus presentation. **(C)** Normalized amplitude spectra for each orientation with the same data in (B), averaged from the 250-500 ms post-stimulus time window. Insert: The normalized mean NBG amplitude of each orientation in polar form of the same data, with the gray arrow indicating the vector mean. **(D)** Normalized mean NBG amplitude of each orientation for each contrast level in Experiment 2, averaged across all VEP electrodes of subject N2-7. Error bar indicates +/- 2 standard error. Together, (B)-(D) show that NBG displays an orientation selectivity to 90°.

Next, orientation tuning strength and the preferred orientation of each electrode was quantified as the angle and magnitude of the orientation selectivity vector (i.e. the mean vector calculated by mean NBG amplitude across orientations). As shown in Figure S2A, the orientation selectivity vector of each VEP electrode and the direction of the orientation selectivity vector summation indicate the general NBG tuning preference around 90°. This general NBG tuning preference is preserved across contrast levels (see Figure S2B-D). Across all electrodes, 53.23% showed a 90° preference (defined as the 45° range with 90° as the center, i.e. 67.5° to 112.5°, same in the following), with 22.58% showing 0° and 24.19% showing 45° or 135° preference. We further employed circular mixed-effect models to evaluate the effect of the contrast level on preferred orientation. The preferred orientation was consistent across contrast levels, as the 95% highest posterior density intervals (HPD) of the circular means for the three conditions overlap (see Table 4). Orientation tuning strength was also consistent across contrast levels (linear mixed effect model, *p* > 0.05). These findings show a striking consistency with prior LFP observations in non-human primates (17), as noted above, and more generally with neuroimaging observations of cardinal biases in human early visual cortex (28).

**Table 4.** Posterior estimates of the preferred orientation (circular mean) for each contrast condition in Experiment 2. s.d. = standard deviation; LB HPD = lower bound of 95% highest posterior density interval; UB HPD = upper bound of 95% highest posterior density interval.

| Condition | Orientation |  |  |  |  |
| --- | --- | --- | --- | --- | --- |
|  | Mode | Mean | s.d. | LB HPD | UB HPD |
| 20% contrast | 83.8° | 85.1° | 13.6° | 57.9° | 111.7° |
| 50% contrast | 73.3° | 76.6° | 12.7° | 50.1° | 100.7° |
| 100% contrast | 93.9° | 92.9° | 12.5° | 67.7° | 117.4° |

In addition to orientation, another key feature encoded in early visual cortex is color. As noted above, prior observations in the non-human primate have reported NBG displays chromatic tuning for long wavelength colors (i.e. red / orange) (18). Recently, we reported that this color tuning of NBG can also be observed in human visual cortex (2), which was later replicated with non-invasive measurements (35). It is therefore of interest to understand how these biases in orientation and color interact to modulate population responses, in particular NBG activity. How these key features of visual cortex selectivity impact NBG activity patterns are critical for understanding its origin and functional significance. Exploring this interaction is further motivated by recent reports showing large scale organization of single-cell joint orientation and color tuning in early visual cortex (25, 36). Prior to examining how color tuning modulates the orientation tuning documented above, we sought to first replicate recent observations of NBG tuning to uniform color stimuli.

### NBG Displays Selectivity to Uniform Color

To confirm NBG color tuning, and ensure it can be observed with penetrating sEEG depth recordings, we replicated the visual color task from Bartoli, *et al*. (2) in the current study. In Experiment 3 (uniform color task), five subjects (N1-3, N6, N7) were presented full screen uniform colors (9 colors spanning long to short wavelengths, 500 ms duration, see Methods).

Raw voltage and bandpass NBG voltage traces for example color trials from an sEEG subject (N6) are shown in Figure 4A. Consistent with previous findings (2), stimulus changes in the raw potential are small in magnitude and difficult to discern. However, when viewing the average spectrograms of the same electrode, clear NBG responses can be observed for red/orange (i.e. long wavelength) and to a lesser extent blue (i.e. short wavelength) hues (Figure 4B). These findings are consistent with observations of ‘end spectral’ biases in early visual cortex (30), whereby greater neural responses are observed for long (red) and short (blue) wavelengths. Averaging NBG amplitude across all subjects further confirmed the preference for red/orange colors (Figure 4C). Strikingly, unlike our prior observations, we found the mean amplitude of induced NBG to green was far higher than that of blue at the group level, in part violating an expected end-spectral tuning profile. Non-parametric testing confirmed a significant modulation of NBG amplitude by color (Friedman test of differences: χ^2^(8) = 29.56; p<0.01). Post-hoc permutation testing confirmed that the NBG amplitude values for red and orange stimuli were all significantly higher than expected by chance (*p-values* < 0.01). Applying permutation testing for each VEP electrode, 100% of all electrodes have significant NBG responses in either red or orange, while 46.9% in blue 1 and 53.1% in green 2 (see Method). Across all VEP electrodes, the preferred color was predominately for the red category (red or orange, 82.8%), followed by the green category (green 1 or green 2, 15.6%) and blue category (blue 1 and blue 2, 1.6%).

**Figure 4.**
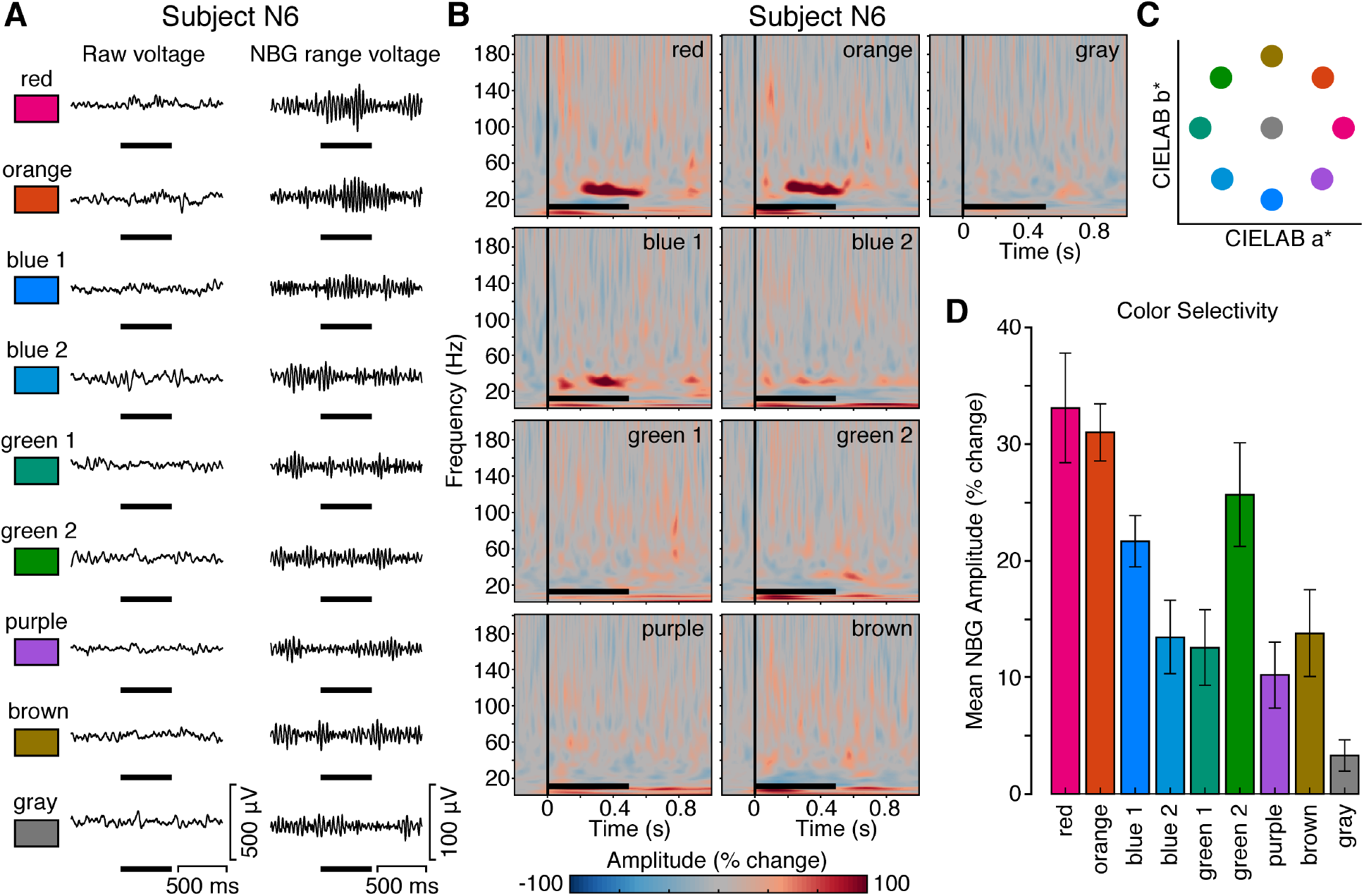
Experiment 3: Spectral Response to Color Stimuli. **(A)** Example single trial voltage response (left: raw, right: bandpass NBG range; from subject N6) for each color (except target color white) in Experiment 3. Full screen uniform colors were presented for 500 ms (black horizontal line), with a random ISI of 1.5-2.0 s. **(B)** Spectrogram for each color from all trials from the same electrode used in (A). Color maps indicate percentage change in amplitude relative to the pre-stimulus period (−500 to 0 ms). Black horizontal line indicates stimulus presentation. Color location in CIE L*a*b* space (lightness plane L = 60) is shown in the last column. **(C)** Group average mean NBG amplitude response across colors (from subject N1-3, N6, and N7; 250-500 ms time window; error bar indicates +/- 1 standard error). Unlike the example electrode shown in (A) and (B), green 2 has higher NBG response compared to blue 1 in the group level, suggesting different subsets of electrodes in color preference.

Related to these observations, it has recently been demonstrated that orientation and color tuning in early visual cortex are more jointly represented than previously appreciated (25). Interestingly, these findings suggest a strong representation of red and blue preferring neurons with strong orientation tuning, providing a potential basis to the stimulus dependence of NBG to both grating and chromatic stimuli (21). As noted above, recent extensions of these findings suggest this strong representation of red/blue is reduced when moving through visual areas V1-V4, such that green becomes more represented and color selective neurons show more spatial clustering (36). Given these joint biases in orientation and color, we would expect interaction effects over NBG modulations when employing stimuli driving both stimulus features. To test this, we examined NBG response to colored grating stimuli.

### NBG Displays Color Modulation of Orientation Tuning

To explore the joint orientation-color tuning of NBG, four subjects (N2-3,N6-7) were presented with full screen chromatic drifting gratings (3 colors: red, green, blue; 8 orientations: 0° to 157.5°, with a 22.5° interval; spatial frequency 1 cycle/degree), for a 500 ms duration (Experiment 4, color grating task, see Methods).

Similar to the responses for uniform color stimuli, chromatic drifting gratings induced relatively modest changes observable in the raw voltage trace (Figure 5A). Interestingly, group averaged spectrograms (Figure 5B) showed NBG responses to each color condition, but with two distinct features. Firstly, responses were far greater, and approximately similar, for red and blue gratings, in contrast to green, as summarized in Figure 5C. Notably, this effect differs from the response to pure color stimuli where responses to green were greater than blue and approximately similar to red/orange on average. This result is potentially consistent with previous evidence of weaker orientation tuning in green-preferring cells (25). As an example of this effect, Figure 5E shows the response for the most green-selective electrode observed to uniform color stimuli, which in turn still shows a consistent red/blue bias for drifting color gratings. Secondly, NBG responses to red and blue are broader in frequency range than that observed for either grating or color stimuli in Experiments 1-3. This broadening of the NBG rhythm further suggests lowered coherence or uniformity in population responses, as well as inter-trial variability, likely a consequence of simultaneously driving joint tuning features. These results indicate that the spatial structure of visual stimuli affects the NBG tuning to fundamental visual features, like color, demonstrating the tight interconnection between orientation and color tuning in human early visual areas.

**Figure 5.**
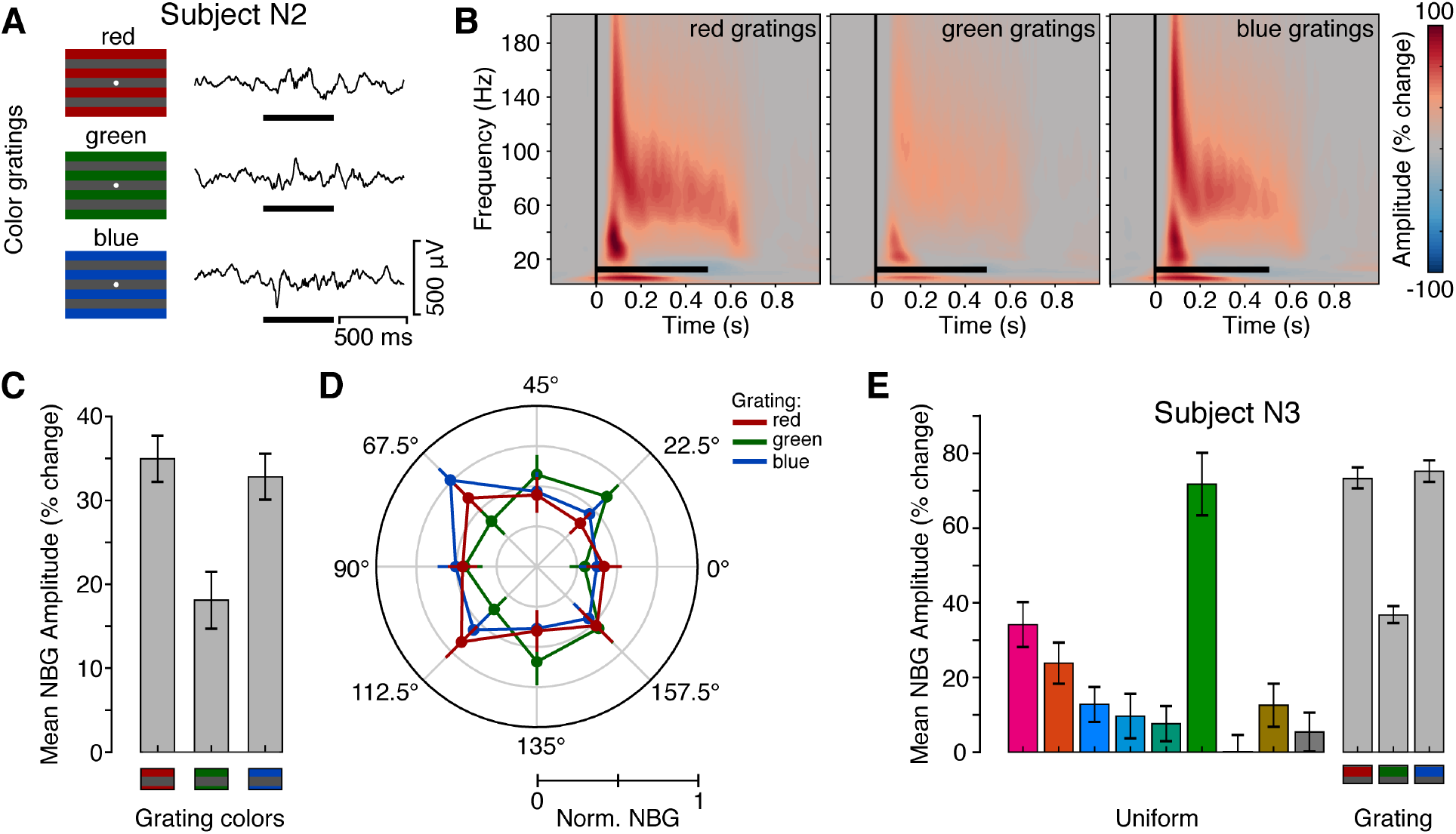
Experiment 4: Spectral Response to Drifting Color Grating Stimuli. **(A)** Example single trial voltage responses (from subject N2) for each grating color in Experiment 4. Grating stimuli were presented for 500 ms (black horizontal line), with a random inter-stimulus interval (ISI) of 1.5-2.0 s. **(B)** Group average spectrograms for each grating color (from subject N2, N3, N6, and N7) in Experiment 4. Color maps indicate percentage change in amplitude relative to the pre-stimulus period (−500 to 0 ms). Black horizontal line indicates stimulus presentation. **(C)** Group average mean NBG amplitude response for color grating task in three color conditions (from subject N2, N3, N6, and N7; 250-500 ms time window; error bar indicates +/- 1 standard error). **(D)** Normalized mean NBG amplitude of each orientation for each color condition (averaged across all VEP electrodes of subject N2, N3, N6, and N7). Error bar indicates +/- 2 standard error. **(E)** Mean NBG amplitude response for uniform color task (Experiment 3) and color grating task (Experiment 4) of a VEP electrode with high green response of N3 (250-500 ms time window; error bar indicates +/- 1 standard error). It shows how the green response was attenuated when color and orientation were jointed driven, in the single electrode level. Together, (B)-(E) shows the interaction between color and orientation in NBG response when these two features were jointly presented.

To further investigate how color and orientation tuning interact, we computed group averaged orientation tuning curves for each grating color. As shown in Figure 5D, orientation tuning is far less pronounced overall, with no color showing a clear bias for 90° like that observed for achromatic gratings. Rather, for blue gratings, NBG orientation tuning is more pronounced for 67.5°. A similar, yet less strongly tuned response, is also seen for red grating stimuli. In contrast to red and blue, green shows a weaker and reciprocal tuning profile, suggesting another domain of dissociation in population responses.

We employed the same quantification of orientation tuning as described for the achromatic drifting gratings. The orientation selectivity vectors of all electrodes across color conditions are less clustered compared to in the achromatic grating task, as shown in Figure S3. Consistent with the orientation tuning curves, the vector summation directions of red and blue are closer to each other compared to green (Figure S3). However, circular mixed-effect model shows the preferred orientations were not significantly different across three color conditions, as the HPD intervals overlap (see Table 5). This may be caused by the relatively small sample size of the current study. In addition, the orientation tuning strength was similar across color conditions (linear mixed effect model, *p* > 0.05). Therefore, grating color not only affected the NBG amplitude response but also the orientation tuning profile of NBG, which does not display the cardinal bias obtained in response to achromatic gratings.

**Table 5.** Posterior estimates of the preferred orientation (circular means) for each color condition in Experiment 4. s.d. = standard deviation; LB HPD = lower bound of 95% highest posterior density interval; UB HPD = upper bound of 95% highest posterior density interval.

|  | **Orientation** |  |  |  |  |
| --- | --- | --- | --- | --- | --- |
| **Condition** | **Mode** | **Mean** | **s.d.** | **LB HPD** | **UB HPD** |
| Red grating | 93.0° | 92.7° | 7.5° | 77.1° | 106.4° |
| Green grating | 74.6° | 74.4° | 12.5° | 49.2° | 98.0° |
| Blue grating | 87.3° | 86.4° | 7.0° | 72.4° | 99.6° |

Together, these findings further support a tight link between the population tuning properties of early visual cortex and the response properties of NBG activity. While NBG reflects known biases for orientation and color, such effects are modulated by their interaction. The differing effects between achromatic and chromatic gratings may be predictive of differential joint tuning biases in population activity, for which recent advances in wide field imaging can examine. As discussed below, these links between ensemble tuning and NBG stimulus dependencies are highly amenable to computational examination and new theoretical views on the functional significance of NBG.

## Discussion

Intracranial recordings (ECoG and sEEG) from human early visual cortex were employed to examine the orientation and color tuning properties of induced narrowband gamma (NBG) oscillations. Consistent with predictions from the non-human primate, we observed both orientation and color tuning of NBG to reflect know biases for cardinal orientations (i.e. 90°), and both end- and mid-spectral color ranges (i.e. red/blue and green). We found that these attributes were modulated when orientation and color were jointly driven by colored-grating stimuli. Red and blue gratings drove greater NBG amplitude responses and showed similar orientation tuning preferences, in contrast to green stimuli. These findings suggest NBG reflects reverberant population activity dependent upon, and sensitivity to, feature selectivity within primary visual cortex. Below we summarize how these findings relate to recent progress in elucidating the large-scale organization of orientation and color tuning in primate visual cortex and how these stimulus dependencies of visual NBG inform theories of its functional significance.

Prior to examining orientation tuning, we replicated previous observations of static grating stimuli reliably inducing NBG activity, whose peak frequency was contrast dependent (2, 16, 31). Whereby, NBG peak frequency increased monotonically with higher contrast levels, as did NBG amplitude. Furthermore, we examined how this relationship was modulated by the use of drifting gratings. As shown in Figure 2, compared to static gratings, drifting gratings produced a uniform increase in NBG peak frequency while still maintaining a monotonic peak frequency increase with contrast level. Grating contrast and velocity modulation of NBG peak frequency has previously been observed with both invasive (14) and non-invasive (37) measurements, adding to other stimulus features which modulate the frequency of visually induced gamma rhythms. It has previously been argued that this stimulus dependence of NBG frequency raises challenges for downstream phase synchronization (16). In addition, while NBG activity is modulated in amplitude and peak frequency by the attributes of grating stimuli, growing evidence shows NBG to be heavily attenuated by complex naturalistic stimuli (2, 11-13). These observations suggest NBG occurs for specific stimulus configurations with high contrast edge/grating like features intersecting the receptive field (21, 38). Given the strong selectivity in early visual cortex for such features, it’s important to examine how stimulus orientation modulates NBG activity.

### Orientation Tuning of NBG

Using drifting grating stimuli, we observed NBG amplitude to be maximal for 90° orientations. This preference for a specific orientation seems unexpected for a population level signal, like NBG, which captures the summed activity of local ensembles with diverse orientation preferences. Consequently, it would be reasonable to predict this summation attenuates any given preference and that NBG amplitude would uniformly increase for all orientations. Importantly, while we did see NBG amplitude increases for all orientations, consistent with this prediction, responses were additionally enhanced for 90° (horizontal) orientations. Interestingly, prior evidence from human neuroimaging studies has shown a cardinal bias in population measures of orientation tuning (28, 29), such that responses are more attenuated for oblique angles. Such a vertical/horizontal orientation bias has also been well known in psychophysical performance (39). Furthermore, work in the non-human primate has also reported a cardinal bias of visual NBG, specifically for 90° orientations (17, 33). More generally, these studies have shown that NBG displays consistent and spatially coherent tuning for a single orientation across proximal recordings sites, despite the underlying spiking activity showing diverse tuning profiles (14, 32, 33). Indicative of this NBG tuning reflecting larger scale population biases, reduced stimulus size (i.e. smaller population activated) greatly reduces the strength and specificity of NBG orientation preference (14). Furthermore, stimulus adaptation to the preferred NBG orientation can produce a transient shift in preference to the orthogonal orientation (14). Together, these observations suggest visual NBG in humans displays a cardinal orientation bias similar to other neural population level measures in humans and non-human primates. However, it is important to note that NBG primarily displayed a preference for one cardinal direction, rather than a general bias for all cardinal directions.

While simple models of coupled inhibitory and excitatory neurons are sufficient to generate rhythmic gamma activity (5), additional circuit features of visual cortex are needed to account for the wide array of tuning properties shown by NBG. In particular, future models will need to account for the multi-scale differences of diverse spiking orientation tuning combining to generate a singular and coherent orientation tuning preference of NBG. As we’ve previously noted, this is a fruitful avenue for developing and testing multi-scale models of population activity (21). However, orientation is not the only key stimulus feature encoded in early visual cortex, color is also strongly represented and has recently been linked to NBG tuning.

### Color Tuning of NBG

Recent work in non-human primates has highlighted NBG also shows color tuning, being particularly enhanced for long wave length (i.e. red/orange), and to a lesser extent, short wave length colors (i.e. blue) (18). We recently replicated this NBG color tuning via invasive recordings in human visual cortex (2), with similar findings subsequently shown via non-invasive measurements (35). Similar to orientation, this color tuning of NBG is consistent with prior neuroimaging reports of an end-spectral bias in population responses within human early visual cortex, particularly V1 (30). Whereby, larger hemodynamic responses were observed for long (i.e. red) and short (i.e. blue) wave length colors. In the present study, we observed a similar influence of color, where NBG responses were largest for red and orange hues. However, unlike our prior work, we observed a larger response to green than blue, which is inconsistent with an end-spectral bias.

What might account for this difference between studies? Regarding stimuli, subjects were presented with the same stimuli via the same apparatus, ruling out this experimental confound. Anatomically, the relative recording coverage of visual areas (V1-4) was also comparable. One key difference is the predominate use of sEEG depth recordings, compared to surface ECoG arrays, in the current study. Penetrating depth electrodes differ in their physical structure, trajectory through the parenchyma and proximity to gray matter, all impacting recorded signals. For example, unlike surface ECoG arrays, sEEG depth electrodes may contact the sulcal wall and fundus, with differing positions relative to the cortical layers. We therefore suspect that the sEEG recordings in the present study allowed for capturing the previously observed end-spectral hue bias, as well as a previously undetected green hue bias. Indeed, there is evidence for mid-spectral bias (i.e. green) in areas V2 and beyond, which may have contributed to our observations (36). While a full retinotopic identification of the visual areas for the current study was not obtained, we employed a probabilistic atlas to infer visual areas, which indicated a sufficient proportion of recordings sites were proximal to V2-4. However, we note that the trajectory of depth electrodes through the occipital lobe allows for capturing activity from multiple visual areas, therefore blurring unique NBG activity from distinct visual areas.

Overall, color tuning of NBG, and its potential variation across visual area, follows a similar relationship to orientation. While red, green and blue selectivity can be observed in early visual cortex, recent findings in wide field imaging have shown a large predominance of red and blue preferring cells in macaque V1 (25, 36). Again, this predominance at the cellular level links to the population tuning properties exhibited by NBG activity. In addition, V1 red/blue predominance changes progressively into downstream visual areas V2-V4, such that the density of green preferring cells increases and that cells sharing the same color preference are more spatially clustered (36). Together these attributes speak to the NBG tuning we observed, which can be considered as a coarse summated sampling of these different populations. One limitation of the present work, motivating futures studies, is to test how closely these changes in cellular color tuning preference and organization map to NBG activity recorded from targeted visual areas (V1-V4). In addition to aiding our interpretation of NBG color tuning, these recent findings also help in understanding why grating-like and uniform color stimuli are particularly effective at driving NBG activity. Specifically, recent findings of joint orientation and color tuning of visual cortex neurons, suggesting overlapping circuits for generating similar NBG dynamics for what otherwise appear as very distinct stimuli. With this unique intersection in mind we examined the influence of joint orientation and color tuning of NBG activity.

### Joint Orientation and Color Tuning in Visual Cortex

Color-grating (red, green, blue) stimuli were used to examine how joint driving of orientation and color influenced the NBG activity we had observed for orientation and color independently. Interestingly, we observed three clear departures from our first set of observations. Firstly, induced NBG was on average broader in its frequency (see Figure 5). Secondly, NBG responses were approximately similar for both red and blue, and comparatively reduced for green – which differs from the clear preferences observed for uniform colors. Thirdly, color-gratings did not drive similar cardinal orientation tuning like achromatic gratings. In contrast, orientation tuning was overall reduced, yet similar in profile for red and blue gratings, while distinct from green orientation tuning. In short, orientation tuning for colored gratings was clearly distinct from that observed with achromatic gratings, and color tuning showed a stronger end-spectral bias than that observed with uniform colors.

What factors can account for these differences? By driving both orientation and color stimulus features one may expect less coherent population level activity (i.e. only specific sub-populations are selectively driven), which is in part supported by the lower overall amplitude of NBG responses to chromatic vs. achromatic gratings. A similar intuition may speak to the broadening of induced NBG frequency. Regarding specific color effects, the reduced responsiveness to green stimuli (when presented as gratings) maybe accounted for by prior evidence of weaker orientation tuning for green preferring cells, compared to red and blue (25). Future studies are needed to examine how cell tuning properties shift over visual areas, for both color preferences and joint color-orientation effects. Based on our data and the interpretation of the observed effects, we would expect reduced evidence for cardinal biases when driving specific color channels.

## Conclusion

NBG oscillations are a striking feature of stimulus induced responses in visual cortex. This high frequency rhythm has been the subject of great interest in theories of visual perception and cognition more broadly. As a periodic modulator of neural excitability, NBG activity is an appealing mechanism for synchronizing neural dynamics. One challenge for such a mechanism is generalization to the rich array of sensory inputs. While classic grating stimuli are highly effective at driving NBG responses, the induced amplitude and frequency of NBG is particularly sensitive to all attributes of such stimuli (e.g. grating size, contrast, spatial frequency, velocity, uniformity). Therefore, even for this idealized stimulus, NBG based synchrony must be adaptive to changes in any of these attributes over time. Given that all of these parameters dynamically change during natural vision, theories suggesting that NBG synchrony supports perception and intra-areal communication must account for continuous changes in magnitude and frequency driven by a host of stimulus dependencies. These challenges grow with consideration of chromatic modulation of NBG properties. In contrast, new hypotheses are emerging, linking NBG occurrence to the presence of specific feature configurations in the visual input able to provoke rhythmic circuit dynamics (19, 21, 27, 38). Computational models will need to account for the diverse parametric relationships between lower level stimulus features and NBG responses (40). Importantly, new theories are needed to help establish general rules of how stimulus composition modulates NBG activity during natural vision (27, 38). Improved computational and experimental links between stimuli, visual circuit properties and the gamma rhythm are critical for elucidating its functional significance.

## Materials and Methods

### Human subjects

Seven subjects (N1-7; 2 males, mean age 37 years, ranging from 22-53 years) were included in this study. All subjects were being monitored with invasive neural recordings (ECoG in N1-2; sEEG, in N3-7) for the potential surgical treatment of refractory epilepsy at Baylor St. Luke’s Medical Center (Houston, Texas, USA). Detailed subject information is reported in Table 1 and Table S1. All subjects participated in this study with both written and verbal consent, completing at least one task in our experimental inventory. Experimental procedures were in accordance with the policy and principles contained in the Declaration of Helsinki and were approved by the Institution Review Board at Baylor College of Medicine (IRB protocol number H-18112). No patients with epileptic foci, anatomical abnormalities or prior surgical resection in posterior regions participated in this study. Experiments were recorded while inter-ictal epileptic discharges were absent in the areas of interest. We note that part of the data recorded from subject N1 (Experiment 3: uniform color) was previously reported in Bartoli, *et al*. (2) under the subject identification code N10. No other datasets from the current study have been previously reported.

### Electrode arrays

Reported data were acquired by ECoG electrode arrays in two subjects (N1-N2) and sEEG depth probes in the remaining 5 subjects (N3-7). The subdural array implanted in subject N1 was custom designed by PMT Corporation (MN, USA) to incorporate a high-density electrode mini grid (4 x 6; 0.5 mm diameter) into a standard macro-ECoG (1 x 8 electrodes with 3 mm diameter) linear array configuration (2, 41). Macro electrodes incorporating the mini grid have an 18 mm center-to-center distance, while other macro electrodes in the array have 10 mm distance. The inter-hemispheric grid (2 x 8) array implanted in subject N2 had dual-sided electrode contacts, with 4 mm diameter and 10 mm center-to-center distance (Ad-Tech Medical Instrument Corporation, WI, USA). Depth sEEG electrodes in subjects N3-7 had a 0.8 mm diameter, with 8 to 16 contacts along the probe with a 3.5 mm center-to-center distance (PMT Corporation, MN, USA). Detailed electrode information for each subject is reported in Table S1 and Figure S1.

### Electrode localization and selection

For each subject, we determined electrode locations by employing the software pipeline iELVis (intracranial Electrode Visualization, (42)). In short, the post-operative CT image was registered to the pre-operative T1 anatomical MRI image using FSL (43). Next, the location of each electrode was identified in the CT-MRI overlay using BioImage Suite (44). For subdural (ECoG) electrodes, coordinates were adjusted for brain shifts by projecting the electrode array to the cortical surface model, reconstructed by the T1 image using Freesurfer (version 5.3; (45)). For the depth sEEG electrodes no location correction was used. For the hybrid mini-macro subdural electrode array in N1, mini-ECoG electrode coordinates were calculated by custom functions combining the macro-ECoG coordinates with the known array geometry.

To identify electrodes in ‘visual’ cortex, we applied an electrode selection criterion previously reported in Bartoli et al. (2019). Briefly, electrodes of interest were those localized within the occipital lobe, with the following anatomical boundaries: 1) dorsal: the parieto-occipital sulcus; 2) ventral: the lingual gyrus; 3) dorso-lateral: the trans-occipital sulcus and posterior aspect of the superior temporal sulcus. Using these criteria, we identified 121 electrodes within the occipital cortex from all 7 subjects (ranging from 8 to 31 electrodes within each subject). We further assigned each electrode to fall within a subdivision of visual occipital areas (V1-4) by mapping the cytoarchitectonic atlas vcAtlas ((46) ; http://vpnl.stanford.edu/vcAtlas) onto each individual cortical surface using Freesurfer. The distribution of electrodes in four subdivisions is reported in Table S1. Occipital electrodes were further selected by means of a functional response criterion (see “Visually responsive electrode identification” below). To visualize electrodes of interest on a common brain, we transformed electrode coordinates into the MNI305 space and represented them as spheres on the Freesurfer fsaverage brain (Figure 1).

### Experimental tasks

All subjects performed at least one of the following visual experiments (see Table 1 for participation details) at the bedside in a quiet patient room. All tasks were presented on an adjustable monitor (1920×1080 resolution, 47.5×26.7 cm screen size, connected to a PC running Windows 10 Pro for subjects N2, N6 and N7 and to an Apple iMac running OSX 10.9.4 for N1, N3-N5) at a viewing distance of 57 cm (such that 1 cm = ∼1 deg visual angle). Tasks were programed using Psychtoolbox-3 functions (v3.0.16) (47) running on MATLAB (R2019a, MathWorks, MA, USA).

### Experiment 1: static grating

Four subjects (N1, N3, N6, and N7) performed the full screen static grating task (see Figure 2), which was previously described in Bartoli et al. (2019) (named as “visual grating task”). During the task, full screen static grayscale grating stimuli were presented for 500 ms, with a random inter-stimulus interval (ISI) between 1500-2000 ms. Stimuli of this task were sine wave gratings at three levels of Michelson contrast (20%, 50% & 100%), and two orientations (0° & 90°). Each block of the static grating task included 105 trials in total, with 90 randomly presented stimulus trials of the 6 conditions (3 contrast levels x 2 orientations, 15 trials for each combination) and 15 target trials presenting 45° gratings at 3 contrast levels (5 trials for each). One block of the static grating task lasted around 4 minutes. Subjects were required to maintain fixation on a white cross at the screen center and respond via button press when a target stimulus occurred. Data from target trials was not included in further analyses.

### Experiment 2: drifting grating

In the drifting grating task, six participants (subject N2-7) were shown full screen moving grayscale grating stimuli (see Figure 2). Grating stimuli had a sine wave spatial frequency of 1 cycle/degree and moved at the speed of 2 degree/sec in two possible directions, orthogonal to the grating orientation. The spatial and temporal frequency settings were chosen to maximize V1/V2 responses (17, 48). Stimuli were presented for 500 ms with a random ISI between 1500-2000 ms. Stimuli in this task reflected three levels of Michelson contrast (20%, 50% & 100%), across eight orientations (0°-157.5° with a 22.5° interval). All eight orientations were covered by two experimental runs: odd blocks contained 0°, 45°, 90° and 135° stimuli, while even blocks contained 22.5°, 67.5°, 112.5° and 157.5° stimuli. Each block of the drifting grating task included 240 randomly presented stimuli from the 12 conditions (3 contrast levels x 4 orientations, 20 trials for each condition), lasting around 10 minutes. N2 and N5 only completed one block of the drifting grating task and do not have data on orientations of 22.5°, 67.5°, 112.5° and 157.5°. Group analyses adjust for this factor. During the task, subjects were required to maintain fixation on a green dot at the center of the screen and press a key when the fixation dot changed size (the fixation dot varied from a 40 pixel diameter to 30 pixel diameter, on one third or the trials). Dot size changed during the 150-450 ms time window after stimulus onset. Data from all trials were included in the analysis.

### Experiment 3: uniform color

Five subjects (N1-3, N6, and N7) performed a full screen uniform color task, which was previously described in Bartoli et al. (2019) (named as “visual color task”). Full screen uniform colors (red, orange, yellow, green1, green2, blue1, blue2, purple, gray and white, see Figure 4 and (2) for more details) were presented for 500 ms with a random ISI between 1500-2000 ms (2). Subjects were required to maintain fixation on a white cross at the center of the screen and press a key when the target color (white) was presented. Each block of the uniform color task randomly presented 140 stimuli (15 trials for each color, 5 trials for target white), lasting around 5 minutes.

### Experiment 4: color grating

In the color grating task, four participants (N2, N3, N6, and N7) were shown full screen chromatic moving grating stimuli (see Figure 5). Grating spatial frequency, velocity and duration parameters were identical to Experiment 2. Gratings were composed of one color (either red, green, blue) against a gray background across eight orientations (0°-157.5° with a 22.5° interval). Colors were selected to closely match the luminance of the gray background and to have a similar color setting with respect to previous studies (19, 25). More details are provided in the section “Control analyses for luminance and chromaticity” below. Similar to the drifting grating task, each block of the color grating task randomly presented 240 stimuli from 12 conditions (3 colors x 4 orientations, either 0°-135° with 45° intervals or 22.5°-157.5° with 45° intervals, 20 trials for each condition), lasting around 10 minutes. Two blocks of different orientation settings were required to show all 8 orientations (N2 only completed one block of the color grating task and does not have data on orientations of 22.5°, 67.5°, 112.5° and 157.5°). Subjects were required to maintain fixation on a white dot at the center of the screen and press a key when the dot changed size (see details from Experiment 2).

### Electrophysiological recording

ECoG and sEEG signals were recorded by a 256 channel BlackRock Cerebus system (BlackRock Microsystems, UT, USA) at 2kHz sampling rate, with a 4th order Butterworth bandpass filter of 0.3-500Hz. ECoG recordings were referenced to an inverted (facing the dura) subdural intracranial electrode, sEEG recordings were referenced to an electrode contact visually determined to be located outside of grey matter. A photodiode sensor recording at 30kHz was attached to the task monitor to synchronize intracranial recordings to the stimulus presentation.

### Data analysis: Preprocessing and spectral decomposition

All signals were processed by custom scripts in MATLAB (R2019a, MathWorks, MA, USA). Raw EEG signals were firstly inspected for line noise, recording artifacts, and interictal epileptic spikes. Electrodes with artifacts and epileptic spikes were excluded from further analysis. Next, each channel was notch filtered (60 Hz and harmonics) and re-referenced to the common average of all selected electrodes. Finally, all re-referenced signals were down sampled to 1kHz and spectrally decomposed using a family of Morlet wavelets, with center frequencies ranging linearly from 2 to 200 Hz in 1 Hz steps (7 cycles).

### Visually responsive electrode identification

Visually responsive electrodes were identified based on the presence of a visual evoked potential (VEP) to grating stimuli, following the definition from a previous study (2). This functional criterion was used to exclude electrodes lacking a robust visual response and at the same time to avoid circularity by focusing on an independent signal feature with respect to the one of interest (i.e. NBG). Electrodes localized within the occipital lobe were defined as “responsive” if: 1) the voltage standard deviation for the response window (0-250 ms) was at least three times greater than the standard deviation for the baseline window (−1000-0 ms); 2) the voltage range for the response window was at least 10 times larger than the standard deviation for the baseline window. The electrodes selected based on the presence of a VEP were 76 % of the electrodes localized to the occipital lobe across all subjects (92 out of 121 electrodes). More than 50% of the visually responsive electrodes were localized in V1 (50 out of 92). Only VEP electrodes were further selected and analyzed (see below). Additional details for each subject are listed in Table S1.

### Statistical analyses

Statistical analyses primarily focused on changes NBG signal properties across task conditions. NBG amplitude was obtained via averaging the magnitude of the Morlet wavelet decomposition result (2 to 200 Hz in 1 Hz steps, 7 cycles) between 20-60 Hz. Amplitude values were normalized to percent change with respect to the pre-stimulus baseline period (500 ms before stimulus onset). As described below, different statistical tests were employed to appropriately evaluate the effects of interest for each analysis. For clarity, they are reported in separate sections below.

### Effect of contrast level and grating motion

To quantify how the amplitude and peak frequency of induced NBG was modulated by the contrast level in Experiments 1 and 2, we computed the log transformed NBG peak amplitude (to correct for skewness) and NBG peak frequency at the single trial level. We employed mixed effects models to evaluate the effect of contrast on NBG amplitude and peak frequency (lme4 library (49) in R (50)). We modeled contrast level as a fixed effect, electrodes and subjects as nested random effects (i.e. full model). To evaluate the impact from the stimulus contrast, we compared the full model to a reduced model with the same random effect structure but no fixed effect (only an intercept term to capture the average). In the results, we report the p-value of the model comparison, assessing the significance of the fixed effect.

To further test whether the amplitude and peak frequency of induced NBG was modulated by the grating motion (i.e. static vs. drifting), we combined the data from experiment 1 and 2 and we modeled both the grating motion and contrast level as fixed effects, using the same nested random effect structure as described above (i.e. full model). By comparing this model with a reduced model (without motion as a fixed effect), we were able to evaluate the importance of grating motion while accounting for contrast. In the results, we report the p-value of the model comparison. The interaction between grating motion and contrast level on mean log NBG amplitude was tested by comparing a model with the additional interaction term to the full model above.

Permutation tests were used for post-hoc comparisons of the grating motion effect at each contrast level. To balance the trial number from each task at a given contrast level, we randomly selected trials from the drifting grating task with the same trial number in the static grating task. Then, the condition labels were shuffled and the difference of NBG peak frequency between the two tasks were calculated for each contrast level. By repeating this process 10,000 times, null distributions of differences at three contrast levels were built. The exact p-value for each contrast level comparison was calculated by the rank of the original difference in the null distribution. To adjust for multiple comparisons, we employed a Bonferroni corrected significance level, reported in the results as p_corr_.

### Orientation tuning

To quantify orientation tuning of NBG amplitude in Experiments 2 and 4, we computed an orientation selectivity vector for each VEP electrode using the following formula:

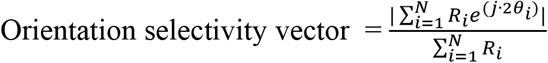

Where *θ*_*i*_ and *R*_*j*_ are the orientations of NBG amplitude at the given orientations. The NBG amplitude values were scaled by the maximum value across all orientations to range between 0-1 for each electrode. Half of the angle of the orientation selectivity vector represents the preferred orientation of the given electrode, while the magnitude of the vector represents the orientation selectivity strength.

We employed circular mixed-effect models (51) to evaluate the contrast effect and color effect on the preferred orientation of VEP electrodes (bpnreg (52) library in R). The contrast level (or color) was modeled as the fixed effect, while subjects were modeled as nested random effects. In the results, we report the 95% highest posterior density intervals (HPD) for each condition. A significant difference would be indicated by non-overlapping HPD of the conditions. In addition, linear mixed-effect models were used to evaluated the contrast effect and color effect on the orientation selectivity strength, similar to what was reported the previous section.

### Color tuning

Previous studies reported that NBG amplitude displays color tuning, especially for reddish-hues. To replicate previous findings, we analyzed the average NBG amplitude values in response to different colors in Experiment 3 using a non-parametric Freidman test of differences as previously described (2). Permutation tests were used to compare the NBG amplitude values of hues to the chance group level (shuffling all the color labels within each subject 10000 times) and electrode level (shuffling the color labels across all trials for each electrode 10000 times). The p-value was calculated by the rank of the original NBG amplitude values in the null distribution.

### Control analyses for contrast level and grating motion effects

To initially assess the influence of grating contrast and motion (static vs. drifting), grating direction of motion and orientation were collapsed. To evaluate whether these two variables would additionally affect how contrast and grating motion modulate NBG amplitude and peak frequency, two control analyses were employed.

First, data from Experiment 2 (drifting grating) was separated by direction and the same modeling as above were employed. No clear differences were observed in models using different sub-datasets (see Table S2). For each grating direction, the effect of contrast was significant for both NBG amplitude and NBG peak frequency (all *p*_*corr*_ < 0.05). Similarly, when testing for static vs. drifting gratings separately for grating direction, both NBG amplitude and NBG peak frequency displayed the same pattern as in the main analysis when the grating directions were collapsed (all *p*_*corr*_ < 0.05).

In the second control analysis, we selected trials at 0° or 90° orientation from the drifting grating task to match the orientations in the static grating task. Results were similar when only trials at 0° or 90° orientation were used (see Table S2). Again, the main effects of contrast and grating motion for both NBG amplitude and NBG peak frequency were significant (all *p*_*corr*_ < 0.05). In summary, NBG peak amplitude and frequency are modulated by contrast levels and by grating motion in a similar fashion when considering different grating orientations and different directions of motion.

### Control analyses for luminance and chromaticity

*CIELxy* coordinates of the stimuli presented in Experiment 3 and 4 were measured using a spectrophotometer (X-Rite i1 Pro, X-Rite, MI, USA). Stimulus luminance in Experiment 3 was greater (∼20 cm/m^2^, see Bartoli et al., 2019 for more details) than Experiment 4 (red: 7.8 cd/m^2^, green: 8.6 cd/m^2^, blue: 9.4 cd/m^2^, gray: 9.3 cd/m^2^). To test if the luminance or other chromatic differences might be affecting the observed results, we evaluated if the mean NBG values obtained in response to each color stimulus in Experiment 3 and Experiment 4 displayed a dependence on CIELxy values measured using multiple regression. We found that the pattern of NBG tuning reported did not depend on luminance values (p=0.12), or the x coordinate (p=0.19) or the y (p=0.29) of CIELxy color space. Therefore, the pattern of color tuning and color-orientation tuning reported in the main analyses cannot be explained by variations in color stimulus luminance settings or by chromatic coordinates in CIELxy space.

## Acknowledgments

This work was supported by NIH grants R01MH106700 and U01NS108923 to S.A.S., R01EY023336 to D.Y., and R01MH116914 to B.L.F. S.A.S is also supported by grants from the McNair Foundation and Dana Foundation.

