## Supplemental Information for "Orientation and color tuning of the human visual gamma rhythm"

#### Supplementary Information

### Figures

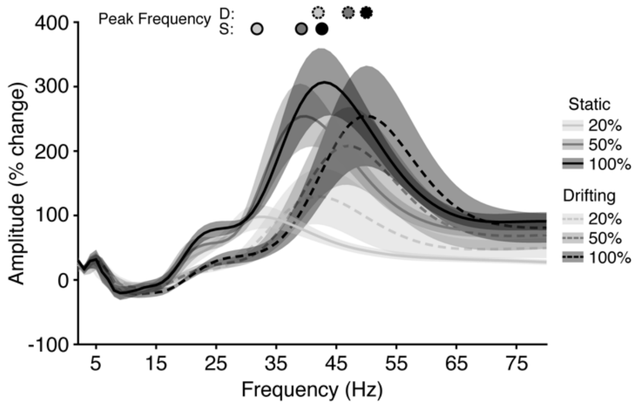


**Figure S1. Group Averaged Spectra for Experiment 1 & 2.** Normalized amplitude spectra were averaged from the 250-500 ms post-stimulus time window. Filled circles indicate peak amplitude frequency. Shading reflects standard error.

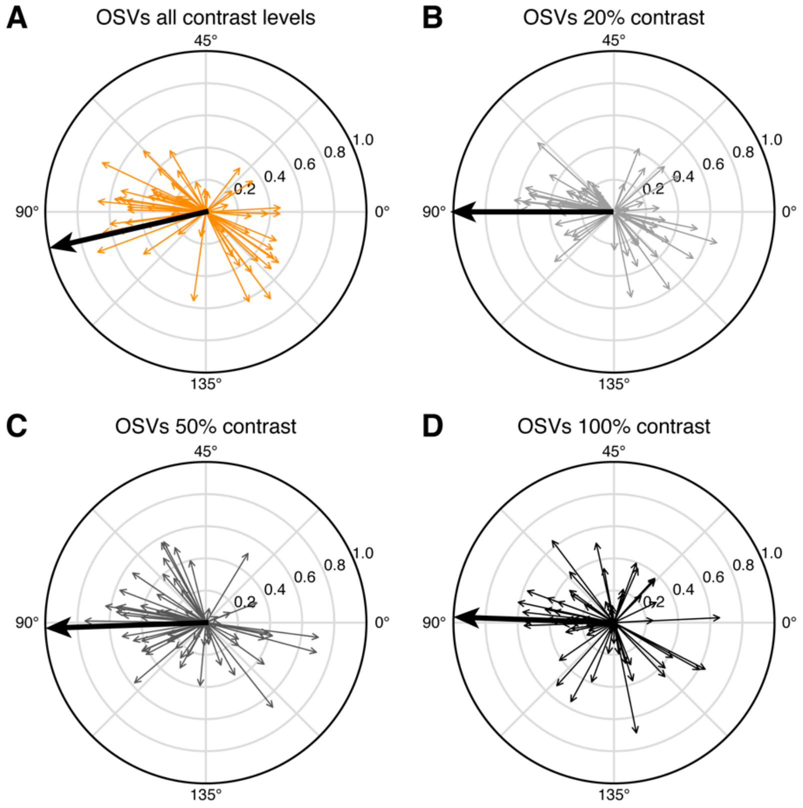


**Figure S2. Orientation Selectivity Vectors (OSVs) of Achromatic Drifting Gratings (Experiment 2).** OSVs of all VEP electrodes from subject N2-7, calculated by NBG amplitudes averaged across three contrast levels **(A)**, at 20% contrast level **(B)**, at 50% contrast level **(C)**, and at 100% contrast level **(D)**. Bold arrows represent the scaled vector summations.

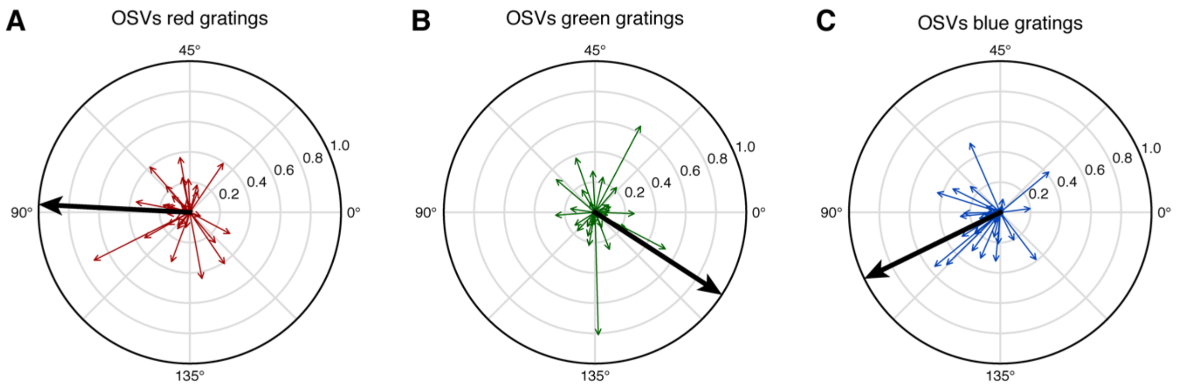


**Figure S3. Orientation Selectivity Vectors (OSVs) of Color Drifting Gratings (Experiment 5).** OSVs of all VEP electrodes from subject N2-3, N6-7, calculated by NBG amplitudes of red gratings **(A)**, of green gratings **(B)**, and of blue gratings **(C)**. Bold arrows represent the scaled vector summations.

### Tables

| Subject code | Hemisphere | Occipital electrodes | VEP electrodes | vcAtlas | | | |
|---|---|---|---|---|---|---|---|
| | | | | V1 | V2 | V3 | V4 |
| N1 | L | 31 | 30 | 25 | 3 | 0 | 0 |
| N2 | B | 12 | 6 | 5 | 1 | 0 | 0 |
| N3 | R | 22 | 14 | 8 | 2 | 1 | 0 |
| N4 | L | 12 | 3 | 0 | 0 | 0 | 1 |
| N5 | L | 26 | 24 | 6 | 1 | 2 | 3 |
| N6 | L | 10 | 7 | 1 | 3 | 0 | 0 |
| N7 | L | 8 | 8 | 5 | 0 | 1 | 0 |
| **Total:** | | **121** | **92** | **50** | **10** | **4** | **4** |

**Table S1. Electrode coverage information.** Hemispheric placement (L = left, R = right, B = bilateral) and count of electrodes in occipital lobe. Electrodes used in the main analyses were selected by means of a functional criterion (displaying a visual evoked potential, VEP). We further assigned the electrodes to different visual areas (v1-v4) based on a probabilistic cytoarchitectonic atlas (vcAtlas, Rosenke*, et al.* (46)) as previously described (2). Note that some electrodes were not assigned to any visual area, as the closest cortical surface location to the electrode coordinate was outside of the atlas boundaries.

| Model components | | Dataset | Fixed parameters (mean + s.e.) | | | |
|---|---|---|---|---|---|---|
| Dependent variable | Fixed effects | | $\beta_{intercept}$ | $\beta_{50\%}$ | $\beta_{100\%}$ | $(\beta_{motion})$ |
| Mean amplitude (% signal change) | Contrast | "Left-down" | 4.5 ± 0.2 | 0.3 ± 0.01 | 0.4 ± 0.01 | |
| | | "Right-up" | 4.6 ± 0.2 | 0.3 ± 0.01 | 0.4 ± 0.01 | |
| | | "0° and 90°" | 4.6 ± 0.2 | 0.3 ± 0.01 | 0.4 ± 0.01 | |
| | Contrast + motion | Static + "Left-down" | 4.5 ± 0.2 | 0.4 ± 0.01 | 0.5 ± 0.01 | 0.03 ± 0.01 |
| | | Static + "Right-up" | 4.5 ± 0.2 | 0.4 ± 0.01 | 0.5 ± 0.01 | 0.07 ± 0.01 |
| | | Static + "0° and 90°" | 4.4 ± 0.2 | 0.4 ± 0.02 | 0.5 ± 0.02 | 0.1 ± 0.01 |
| Peak frequency (Hz) | Contrast | "Left-down" | 38.8 ± 0.4 | 5.4 ± 0.2 | 8.0 ± 0.2 | |
| | | "Right-up" | 38.2 ± 0.4 | 5.2 ± 0.2 | 7.6 ± 0.2 | |
| | | "0° and 90°" | 37.3 ± 0.7 | 5.4 ± 0.3 | 9.0 ± 0.3 | |
| | Contrast + motion | Static + "Left-down" | 33.1 ± 0.3 | 5.3 ± 0.2 | 7.8 ± 0.2 | 5.8 ± 0.2 |
| | | Static + "Right-up" | 33.3 ± 0.3 | 5.2 ± 0.2 | 7.5 ± 0.2 | 5.0 ± 0.2 |
| | | Static + "0° and 90°" | 33.0 ± 0.3 | 5.3 ± 0.2 | 8.3 ± 0.2 | 4.5 ± 0.2 |

**Table S2. Coefficients of fixed parameters in control analyses.** $\beta_{intercept}$ captures the 20% contrast level mean in each model. $\beta_{50\%}$ captures the mean difference between 50% and 20% contrast. $\beta_{100\%}$ captures the mean difference between 100% and 20% contrast. $\beta_{task}$ captures the mean difference between the drifting grating task and static grating task in the complex models. "Left-down", "Right-up" and "0° and 90°" indicate three subsets of data from the drifting grating task.
